# A high-throughput approach to identify reproductive toxicants among environmental chemicals using an *in vivo* evaluation of gametogenesis in budding yeast *Saccharomyces cerevisiae*

**DOI:** 10.1101/2022.01.25.477777

**Authors:** Ravinder Kumar, Ashwini Oke, Beth Rockmill, Matthew de Cruz, Rafael Verduzco, Xavier W. Madeira, Dimitri P. Abrahamsson, Joshua F. Robinson, Patrick Allard, Tracey J. Woodruff, Jennifer C. Fung

## Abstract

**Background:** Environmental chemical exposures are likely making important contributions to current levels of infertility and its increasing incidence. Yet the US produces high volumes of industrial chemicals for which there is limited data on their potential human reproduction toxicity. Current assays typically used in policy and regulatory settings involve costly and time-consuming whole-animal rodent tests which limit the rapidity with which one can assess the thousands of chemicals yet to be tested.

**Objective:** Our aim was to develop a fast and reliable strategy to evaluate a large number of chemicals for reproductive toxicity by developing a high-throughput toxicity assessment using the yeast *S. cerevisiae.*

**Methods:** Yeast are chronically exposed to each environmental chemical at two doses, 30 μM and 100 μM, in a 96-well plate-based format throughout gametogenesis. Non-gametes are removed and chemicals are washed away before gamete viability is measured using absorbance at 600 nm to produce growth curves. The difference in time at half-maximal saturation with and without exposure is used to determine the extent of reproductive toxicity.

**Results:** We validated our assay using bisphenol A (BPA), a well-established mammalian reproductive toxicant. We find that BPA in yeast has similar detrimental effects in meiosis as shown in worms and mammals. Competition assays with BPA analogs reveal that two of out of 19 BPA analogs examined (bisphenol E and 17β-estradiol) show synergistic effects with BPA at doses tested and none show antagonistic effects. Out of 179 additional environmental chemicals, we designated 57 chemicals as reproductively toxic. Finally, by comparing chemicals in our cohort that have been evaluated for reproductive toxicity in mammalian studies, we find a statistically significant association between toxic chemicals in yeast and mammals.

**Conclusion:** We show that a high-throughput assay using yeast may be a useful approach for rapidly and reliably identifying chemicals that pose a reproductive risk.

## Introduction

Infertility is a surprisingly common problem affecting 10-15% of reproductive age couples (Hull et al. 1985). It can stem from a variety of causes including reduced quality and quantity of gametes (both sperm and eggs), physical blockage of the male or female ducts, as well as uterine abnormalities. In the broad class of infertility in which quality and quantity of gametes are reduced, problems of gametogenesis are a main contributor (Hassold and Hunt, 2007). Failure of gametogenesis is primarily due to a breakdown in the ability of chromosomes to divide properly during meiosis, ultimately resulting in gamete aneuploidy. In addition to infertility, gamete aneuploidy will also manifest as an increased incidence of miscarriages in the mother and developmental disabilities in subsequent generations (e.g. Down Syndrome – trisomy 21, Edward’s Syndrome – trisomy 18, Patau Syndrome – trisomy 13) (Nagaoka et al. 2012).

Not all underlying causes for gamete aneuploidy are known, however there is mounting evidence that environmental chemicals can contribute to its incidence (Jorgensen et al 2021; Lea et al 2016; Skakkebæk et al 2012). Very little is known about the reproductive and developmental toxicity of the majority of industrial chemicals in use in the United States, even for those commonly detected in maternal and umbilical cord sera (Panagopoulos et al 2021; Wang et al 2021). There are currently over 86,000 industrial chemicals listed in the U.S. Toxic Substances Control Act (TSCA, 15 U.S.C. §§ 2601 et seq.) chemical inventory, of which approximately 40,000 (47%) are actively manufactured, imported, or used in household or commercial products. The majority of industrial chemicals have never been evaluated for their potential toxicities towards human health and reliable information about reproductive toxicology is particularly scarce (Di Renzo et al 2015).

In humans, an impediment to identifying reproductive toxicants is the prolonged delay between toxicant exposure and the manifestation of reproductive perturbations. Meiosis, a key molecular process leading to gametogenesis, takes place in female fetuses *in utero* and manifestation of the adverse effect is not observed until adulthood. If a woman is exposed to meiotic toxicants as an adult, they will not necessarily affect her own fertility, since a significant part of meiosis has already taken place during her fetal ovarian development. Instead, when a pregnant woman is exposed to a reproductive toxicant, the effects may only be seen when her children attempt to conceive or when her grandchildren are born. Relating chemical exposure in an individual to a fertility reduction in their children, or birth defects in their grandchildren, is epidemiologically challenging due to the need for large cohorts followed over long periods of time with adequate information about exposure during critical developmental periods. As a result, there is a relative paucity of information on human reproductive toxicity, although ongoing and multiple exposures to environmental chemicals has been well documented (reviewed in Hoyer 2001, Younglai et al 2005).

Evaluation of reproductive toxicity is most commonly performed using whole-animal rodent tests. But because in mice, just like in humans, the reproductive effect of chemical exposure in a pregnant female may only become apparent one or two generations later, these experiments are costly, time consuming, and require a large number of animals for transgenerational studies, thus greatly restricting the number of chemicals that can realistically be tested. At the same time, the use of mammals for chemical testing raises ethical considerations, spurring an interest in looking for new approaches that would avoid the use of mammals as a primary testing method. Due to ongoing chemical exposures with little data on reproductive toxicity, there is an urgent need to develop approaches for more rapid testing.

Recently, there is growing interest in the regulatory community to develop a more holistic assessment of toxicity by incorporating multiple lines of evidence through computational modeling (Thomas et al 2019) that integrates *in vitro* assays that assess related biological pathways (Blaauboer 2008; Kavlock and Dix 2010), quantitative structure-activity relationships (QSARs) that map molecular structural features of chemicals to their physical, chemical, or biological properties (Gini 2018; Liu et al 2017; Low et al 2011) and *in vivo* assays that do not utilize mammalian models but instead employ alternative organisms (Allard et al 2013; Cornet et al; 2017; Ferreira and Allard 2015; McGrath and Li; 2008). These types of analyses rely on high-throughput assays to provide the data that can augment toxicity assessments. But the challenges of assessing reproductive toxicology in mammals prohibits the acquisition of quantitative toxicology data on the large scale needed for these data-intensive approaches.

One system that would be amenable for high-throughput discovery of reproductive toxicants is the budding yeast, *Saccharomyces cerevisiae.* Yeast has long been a major workhorse of eukaryotic molecular biology, and in addition to the leading role it has played in our understanding of such common core processes as transcription, chromatin, DNA replication, and the cell cycle, yeast is one of the most studied organisms for gametogenesis (Roeder 1995). The use of this organism thus leverages an extensively developed trove of molecular and genetic information. Moreover, conservation is remarkably high between yeast and humans with ~60% of yeast genes having human homologs and 87% of yeast protein domains being present in the human proteome (Peterson et al 2013). In terms of assaying meiotic toxicants, yeast has the added benefit that, as a single cell system, it lacks a reproductive tract, thus gametogenesis can be evaluated directly. Yeast is also easily induced to undergo gametogenesis by a simple exchange of growth medium to one deprived of nutrients to which chemicals can be introduced at the same time. Yeast has also proven to be useful in high-throughput experiments because of its rapid growth in liquid media that allows experiments to be conducted in multi-well plates.

Yeast-based proliferative high-throughput screens (HTS) have been successfully used to identify compounds that target conserved proteins. This has been exemplified by the ability of several high use drugs—statins, omeprazole, tacrolimus, bortezomib and methotrexate – to hit the same highly conserved targets and elicit the same responses in yeast as in humans (Armour and Lum 2005; Kachroo et al. 2005; Tardiff et al. 2012, 2013).

In this study, we developed a yeast-based HTS to evaluate reproductive toxicity. We first evaluated our assay’s ability to detect reproductive toxicants by testing BPA, a plastic precursor that has been detected in human tissues and that has known adverse effects on mammalian reproduction (reviewed in Modal et al. 2021, Vandenberg et al. 2019). Due to regulatory and marketing activities to reduce BPA exposures, use of BPA has decreased, but this has led to an increase in substitution with BPA analogs (Catenza et al. 2021) for which the potential for toxicity is not well characterized (Pelch et al. 2017; 2019). Given the structural similarities of these analogs to BPA, and the concomitant potential for health effects, we evaluated the toxic effect of BPA analogs on gametogenesis by examining 19 BPA-related compounds. Moreover, in view of the potential concurrent exposures to BPA and its alternatives, we performed a series of competitive assays to elucidate whether these chemical mixtures act additively, synergistically or antagonistically. Finally, we expanded our test system to evaluate the potential reproductive toxicity of 179 additional chemicals that we identified as high priority (Abrahamsson et al. 2022). We also compare the findings from our *in vivo* yeast assay to reproductive toxicity in mammalian studies

## Methods

### Yeast strains and growth media

Yeast strains were constructed in a diploid BR1919-8B background which has high meiotic efficiency (Rockmill and Roeder 1998). Alleles are homozygous in diploid strains except where explicitly noted. The chemical sensitized strain is SRK007 *(MATa/MAT**α**ADE2:ade2-1*, *leu2-3,122 his4-260 ura3-1 thr1-4 lys2 trp1-289 pdr1::KANMX pdr3::NAT*). In order to obtain genetic control strains for evaluating assay performance, mutations known to affect meiosis were added by crossing SRK037 (MAT*α pdr1D pdr3Δ*) to JCF28 *(msh4::URA3)* and JCF249 *(spo11:ADE2)* to generate diploids SRK041 *(MATa/MAT**α** pdr1D pdr3D msh4::URA3)* and SRK040 *(MATa/MAT***α***pdr1.Δ pdr3D spo11::ADE2*). Yeast strains are kept in 50% glycerol frozen stocks and grown on YPD media supplemented with uracil and adenine (1% yeast extract, 2% bactopeptone, 2% dextrose, 0.9% adenine and 0.2% uracil) for proliferative growth. To induce gametogenesis, T-SPO media (1% potassium acetate, 0.1% yeast extract and 0.05% dextrose) is used.

### Chemicals

Table S1 lists the chemicals used in this study with the associated catalog numbers, manufacturer, CAS number, barcode and % purity, usage and chemical class. Table S2 lists the associated chemical structures as determined by ClassyFire (Djoumbou et al. 2016) for each chemical. Seven replicate 100 mM stocks were made in 100% DMSO (Sigma Aldrich, St. Louis, MO) and were stored in 0.5 ml aliquots in 1.10ml polypropylene, screw cap, barcoded tubes with internal threads (Micronic America, Aston, PA) at −80°C. Chemicals were thawed for use and diluted to exposure concentrations while keeping the final DMSO concentration at 0.1%. On occasion, 10 mM stocks were made due to limited chemical availability or solubility.

### Yeast reproductive toxicity assay

#### Cell preparation

The SRK007 yeast strain is freshly streaked from frozen stock and grown on YPD plates. A saturated culture is generated by inoculating a colony into 1 ml of YPD and grown for 24 hours at 30°C on a shaker set at 230 rpm. After 24 hours, cells are pelleted and washed three times with T-SPO media before resuspension into 1 ml of T-SPO. Cells are diluted into 50 ml of T-SPO to a final OD600 of 0.25. 100 μl of cell suspension is transferred into each well of a 96 deep-well plate using a Liquidator-96 (Mettler-Toledo Rainin, LLC, Columbus, OH) into which 400 μl chemicals and sporulation media have been dispensed such that the final doses are 30 μM or 100 μM in a 0.5 ml volume. The plate is covered with a Breathe Easier membrane (Sigma-Aldrich, St. Louis, MO) before incubation in a Multitron HT shaker (Infors AG, Basel, CHE) at 30°C, 950 rpm for 72 hours.

#### Chemical dispensation

Chemicals are dispensed just prior to cell inoculation into sterile 96-deep well plates with pyramidal bottoms. Only the inner 60 wells are used. To prevent evaporation, the outer 36 wells are filled with 0.5 ml of sterile water. 100 mM chemical stocks are first thawed and vortexed before diluting into two working solutions of 37.5 μM and 125 μM from which 400 μl is added to each well. Once the cells are added, three replicates at the final dose concentrations of 30 and 100 μM in 0.1% DMSO are made. For each plate, four replicates of the negative control 0.1% DMSO and positive control 0.5 μM Latrunculin B (LatB) are also included. Different chemicals were randomly positioned for each plate.

#### Growth curve measurement

On completion of incubation, cells are pelleted in a tabletop centrifuge at 1000 x g for 5 minutes. Cells are washed and pelleted three times with 300 μl sterile water. Cells are resuspended in 100T zymolyase (31.25 μg/ml of zymolyase 100T and 10 mM DTT) and incubated at 30 °C for 3 hours. After each hour, cells are vigorously mixed in a Mixmate (Eppendorf, Hamburg DE) at 1000 rpm for 2 minutes. After incubation, cells are washed three times with 300 μl of sterile water. Cells are then resuspended in 1 ml YPD and vigorously mixed in the Mixmate for 10 minutes. 10 μl of the cell suspension is added to 90 μl of YPD in each of the inner 60 wells of a 96-well imaging plate (Corning #3631, Corning NY). The outer wells are filled with 100 ul of YPD to limit evaporation, the plate is covered and sealed with tape on all sides before placing in the Tecan M200 plate reader (Tecan, Männedorf, CHE) with the following settings (30°C, 432 orbital shaking,1 mm amplitude). Readings are recorded every 30 minutes for 35-45 hours using Tecan’s iControl software.

#### Δt_Hmax_ *calculation*

OD values across all timepoints are used to fit a logistic growth curve using R package drc (Ritz et al. 2015) with the following parameters (fct=l4(fixed=c(NA,NA,1,NA)) for slope, start and end of curve and intercept. Time at half-max (t_Hmax_) is calculated as time to reach OD 0.5 on the fitted growth curve. For each chemical, the shift in t_Hmax_, Δt_Hmax_ is calculated as the difference between a chemical’s t_Hmax_ and the average t_Hmax_ of the 0.1% DMSO wells on that plate. For wells that do not reach saturation, the t_Hmax_ value is capped at 46 hr (32 hr for mitotic experiments). Wells with poor fits are flagged and removed from the data set.

#### Meiotic characterization

Chromosome spreads (Rockmill 2009) were prepared at 19 hours and 22 hours after meiotic induction and imaged on a Deltavision (GE Healthcare) fluorescence microscope. Spreads were stained with anti-Zip1 antibodies to highlight the synaptonemal complex and anti-Rap1 antibodies to highlight the ends of the chromosomes. Each chromosome spread was evaluated for the state of synapsis progression (MacQueen and Roeder 2009). The frequency of cells that progress beyond meiosis I was calculated from counting the number of nuclei 3 days after cells were induced to undergo meiosis and then fixed with 70% ethanol and stained with DAPI to highlight the number of nuclei. Gamete viability was determined by manually dissecting 10 mM zymolyase-digested 4-spore viable tetrads onto YPD plates (Guthrie and Fink 1991). Recombination was measured in centiMorgans (cM) for *HIS4-LEU2* and *LEU2-MAT* intervals based on the number of parental (P), nonparental ditype (NPD) and tetratype (T) combination of genetic markers (Perkins, 1949).

#### BPA and BPA Substitute Competitive Assays

For those BPA alternatives that at 15 μM or 30 μM shifted the t_Hmax_ without affecting either slope or saturation, we determined combination effects using the Loewe additivity model (Chou and Talalay 1984; Gaikani et al. 2021; Loewe 1953). Δt_Hmax_ was calculated for BPA and a BPA alternative individually at X μM, 2X μM and mixed at X μM BPA substitute + X μM BPA doses where X could be 15 or 30 μM. A synergistic effect is concluded if Δt_Hmax_ for the X μM BPA substitute + X μM BPA doses is significantly greater than for the 2X μM doses of either BPA alone or its substitute alone. Antagonistic effects are concluded when Δt_Hmax_ for X μM BPA alternative + X μM BPA doses is less than for both 2X μM BPA or 2X μM BPA substitute.

## Results

### Yeast high-throughput screen for reproductive toxicants

To rapidly assess a large number of chemicals for reproductive toxicity, we developed a 96-well plate high-throughput screen (HTS) based on detecting gamete viability in budding yeast (**Figure 1**). Because gametes are haploid, and each chromosome carries essential genes, any failure of meiotic chromosome segregation leading to chromosome loss would produce inviable gametes. Since yeast can proliferate in either the diploid or haploid state, the level of gamete viability in yeast is an easily measurable indicator of meiotic success as it relies only on absorbance measurements to assess the proliferative growth of viable gametes, making it more suitable for high throughput screening. Compounds detrimental to meiosis will cause a shift in the gamete growth curve to the right relative to the vehicle control due to the decreased number of viable gametes (**Figure 1**) which delays the appearance of visible growth. In this assay, the chemical being tested is only applied while the cells are undergoing meiosis, and is then extensively washed away so that growth curves are measured in the absence of the chemical (see Methods). The shift in time observed at half maximum of the growth curve (Δt_Hmax_) reflects the number of viable cells present in the sample at the start of the growth phase, and therefore the extent of toxicity. To reduce the well-known resistance of yeast to exogenous chemicals, we constructed a *pdr1Δ pdr3Δ* double-mutant strain that codes for transcription factors needed for the MDR class drug efflux pumps (Delaveau et al. 1994). The *pdr1Δ pdr3Δ* double-mutant strain has been used effectively in a HTS for drugs that affect neurodegenerative disease (Tardiff et al. 2012) and we found that it does not significantly affect gamete viability (**Figure S1A**). A key aspect of the assay is the removal of any diploid cells that fail to enter meiosis that would otherwise confound the gamete viability assessment after the cells are reintroduced to proliferative growth. Diploids that do not undergo meiosis are removed by enzymatic digestion with zymolyase, which digests the cell wall of vegetative cells, but which is much less effective at digesting the spore wall that protects the gametes (Wolska-Mitaszko et al. 1981). Once diploids are removed and the chemicals washed away, gamete viability can be assessed using absorbance at OD_600_ in a microplate reader by measuring resumption of proliferative growth after exchange into YPD medium. Proliferation is monitored until the growth curve reaches saturation in order to accurately calculate the Δt_Hmax_ and to detect if sustained damage occurred that affects mitotic/proliferative growth (e.g. losses to mitochondrial function) which is evident by both a change in slope in the growth curve and a lowered saturation level (**Figure 1**, arrowhead). Because these growth curves are obtained after the chemicals have been fully washed away, any delay in growth is the result of earlier defects sustained during the meiotic process, such as loss of essential genes due to chromosome mis-segregation.

**Figure 1.**
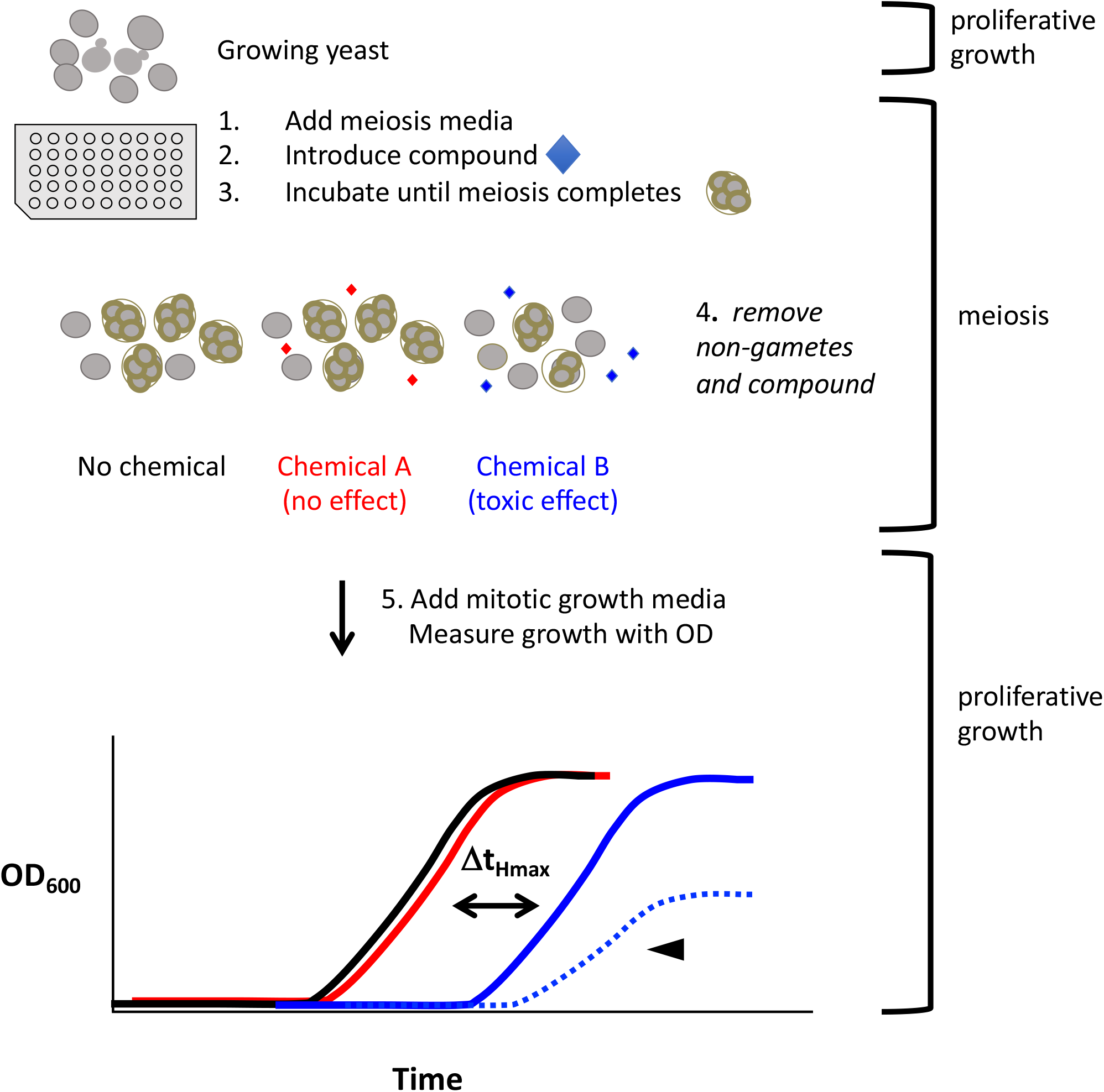
Yeast HTS for reproductive toxicity. Chemicals (blue diamonds) are introduced to yeast in meiosis media used to initiate gametogenesis. After gametogenesis completes with the formation of a four-gamete tetrad, non-gametes are removed by prolonged zymolyase digestion which digests the cell wall of diploid cells leaving them open to osmotic shock. Gametes survive due to the spore wall that is unaffected by the zymolyase digestion. Remaining chemicals are removed by several wash steps before the gametes are reintroduced to proliferative growth. Growth curves are obtained by measurement at OD600 in a plate reader. Any toxic chemical that reduces the number of gametes or decreases gamete viability will cause the growth curve to shift to the right relative to the no chemical control. The measured shift in time at Δt_Hmax_ reflects the extent of toxicity. Any growth curves that show a change in slope and/or lowered plateau reflect acute toxicity and t_Hmax_ is capped at 46 hours.

### Validation of screen using meiotic mutants and bisphenol A (BPA)

As an initial validation of our approach for using growth of meiotic products to detect meiotic defects, we examined two well-characterized yeast meiotic mutants, *spo11Δ* (Keeney et al. 1997) and *msh4Δ* (Ross-Macdonald and Roeder 1994), both of which are recombination mutants with known loss of gamete viability (<1% (Rockmill et al. 2013) and 43% (Chen et al. 2008), respectively). As seen in **Figure 2A**, both mutants shift the growth curves rightward to an extent compatible with their known gamete viability (**Figure 2B**) suggesting that the assay accurately reflects meiotic perturbations in viable gamete number. To determine whether our assay is sensitive enough for HTS applications, we calculated a Z’ value (Zhang et al. 1999), a robust measure of separation between hits and non-hits in a screening experiment. A Z’ of 0.753 was determined from the signal dynamic range and data variation from both the negative control (0.1% DMSO) and the positive control (0.5 μM Latrunculin B (LatB)). Z’ values between 0.5 to 1.0 indicate a high quality HTS. Latrunculin B (LatB), a highly specific inhibitor of the actin cytoskeleton, was used as a positive control since it is known to disrupt the cytoskeletal elements needed for telomere-led chromosome motion in budding yeast essential to prophase I of meiosis (Koszul et al. 2008). Our negative control – 0.1% DMSO was selected to solubilize the chemicals since it had no effect on meiosis up to 1% DMSO (**Figure S2B**) and can solubilize a wide variety of otherwise poorly soluble polar and nonpolar molecules. Similarly, no toxicity was observed for acetone, acetonitrile, methanol and toluene. Among potential solvents tested, ethanol was detrimental to meiosis, as ethanol can be used as a carbon source thus preventing meiotic entry.

**Figure 2.**
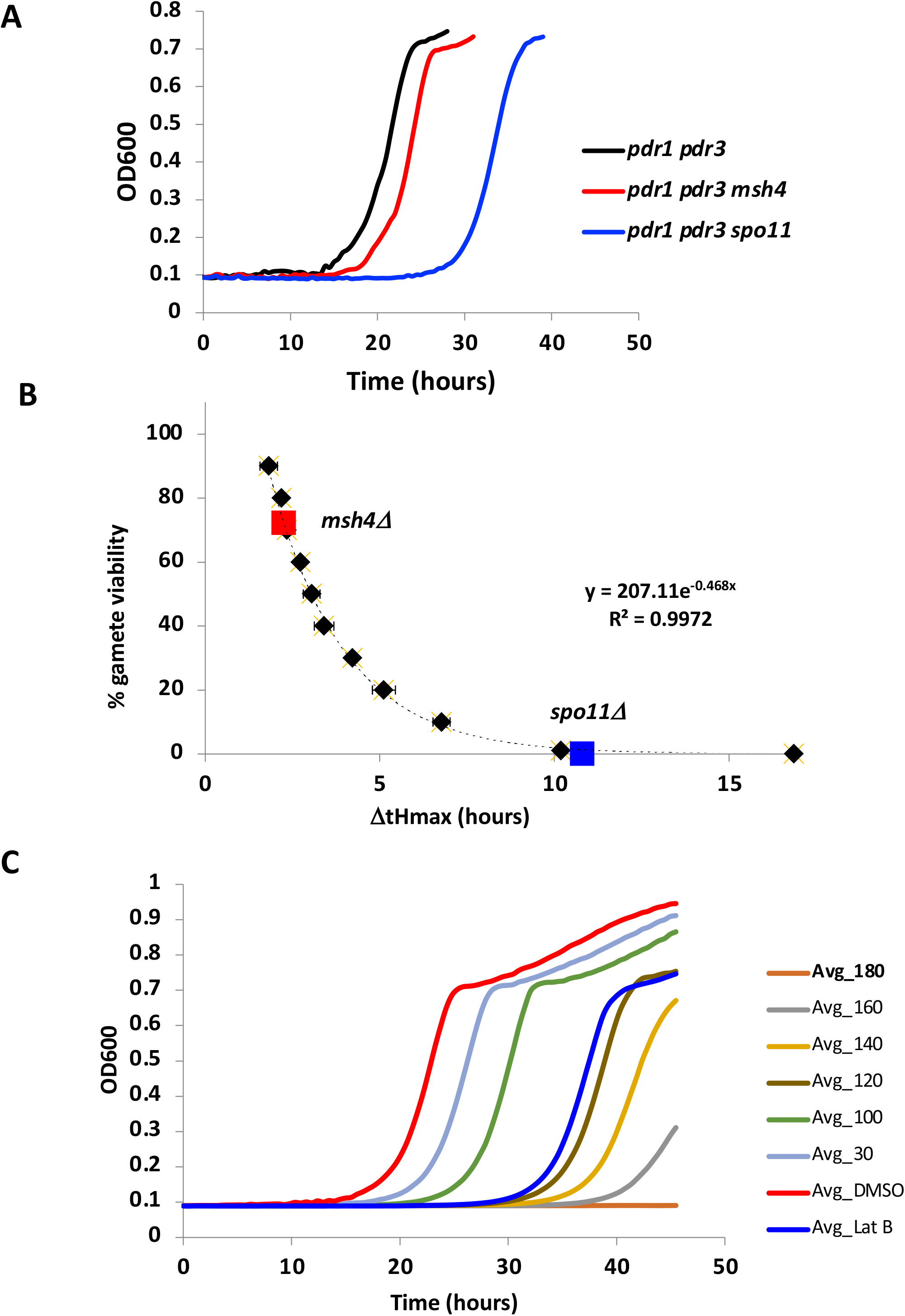
Validation of screen using meiotic mutants and known reproductive toxicant BPA. A) Growth curves for meiotic mutants *msh4* and *spo11* B) Mutant Δt_Hmax_ mapped onto standard curve calculated from a dilution series of sporulated cells (black diamonds). Formula for curve fit used to convert Δt_Hmax_ to % gamete viability C) Dose response curves for BPA at several increasing concentrations shown in μM. The experiments were performed in triplicate and averaged values are shown. DMSO – 0.1% (negative control). LatB – 0.5 μM (positive control).

To ask if our assay detects reproductive toxicants, we tested bisphenol A (BPA), a well-studied chemical with known adverse effects on mammalian reproduction. BPA (Hunt et al. 2003; Susiarjo et al. 2007; Vrooman et al. 2015) is a plastic precursor that has been found extensively in humans (Gerona et al. 2016; Li et al. 2022; Vandenberg et al. 2007) due to its widespread use in food and beverage containers, thermal paper, toys, electronics, medical equipment and water pipes (reviewed in Catenza et al. 2020). We tested BPA’s dose response at 0, 30, 100, 120, 140, 160, 180 μM. As shown in **Figure 2C**, our assay exhibits sensitivity to BPA, showing greater toxicity (i.e. larger shifts in Δt_Hmax_) with greater dose.

Prior studies of both mammals and worms have shown that BPA disrupts meiosis (Susiarjo et al., 2007; Allard et al., 2010). During prophase I oogenesis in mice, Susiarjo et al. (2007) observed both unsynapsed chromosomes and higher recombination that resulted in an increase in aneuploidy. In nematodes, Allard et al. (2010) also found unsynapsed chromosomes and a delay in double-stranded break processing during meiosis resulting in fewer eggs and higher embryonic lethality. In yeast, we see a similar perturbation in meiosis during prophase I at 19 hours (**Figure 3A**) and 22 hours (**Figure 3B**) after meiotic induction. This is manifested both as a delay in chromosome synapsis progression and by the unexpected appearance of the polycomplex, an abnormal aggregation of the synapsis protein Zip1 previously shown to accompany problems in chromosome synapsis (Sym and Roeder 1995). Further microscopic evaluation of BPA effects at 30 and 100 μM in yeast reveals that the overall frequency of cells progressing through meiosis is reduced at both doses (**Figure 3C**), however gamete viability is only significantly perturbed at 100 μM BPA (**Figure 3D**). Changes to recombination can often lead to loss of gamete viability. To determine if recombination is affected by BPA, we measured recombination in two genetic intervals, *HIS4-LEU2* and *LEU2-MAT*. Although recombination was not affected at 30 μM, we observed reduced recombination at 100 μM BPA as compared to 0 μM BPA *(HIS4-LEU2:* 23.6 cM to 12.0 cM; *LEU2-MAT:* 32.3 cM to 19.1 cM) (**Figure 3E**). Together these results show that BPA in yeast causes defects in meiotic prophase I as observed in both mammals and worms, suggesting that BPA affects the same conserved mechanism in these diverse organisms.

**Figure 3.**
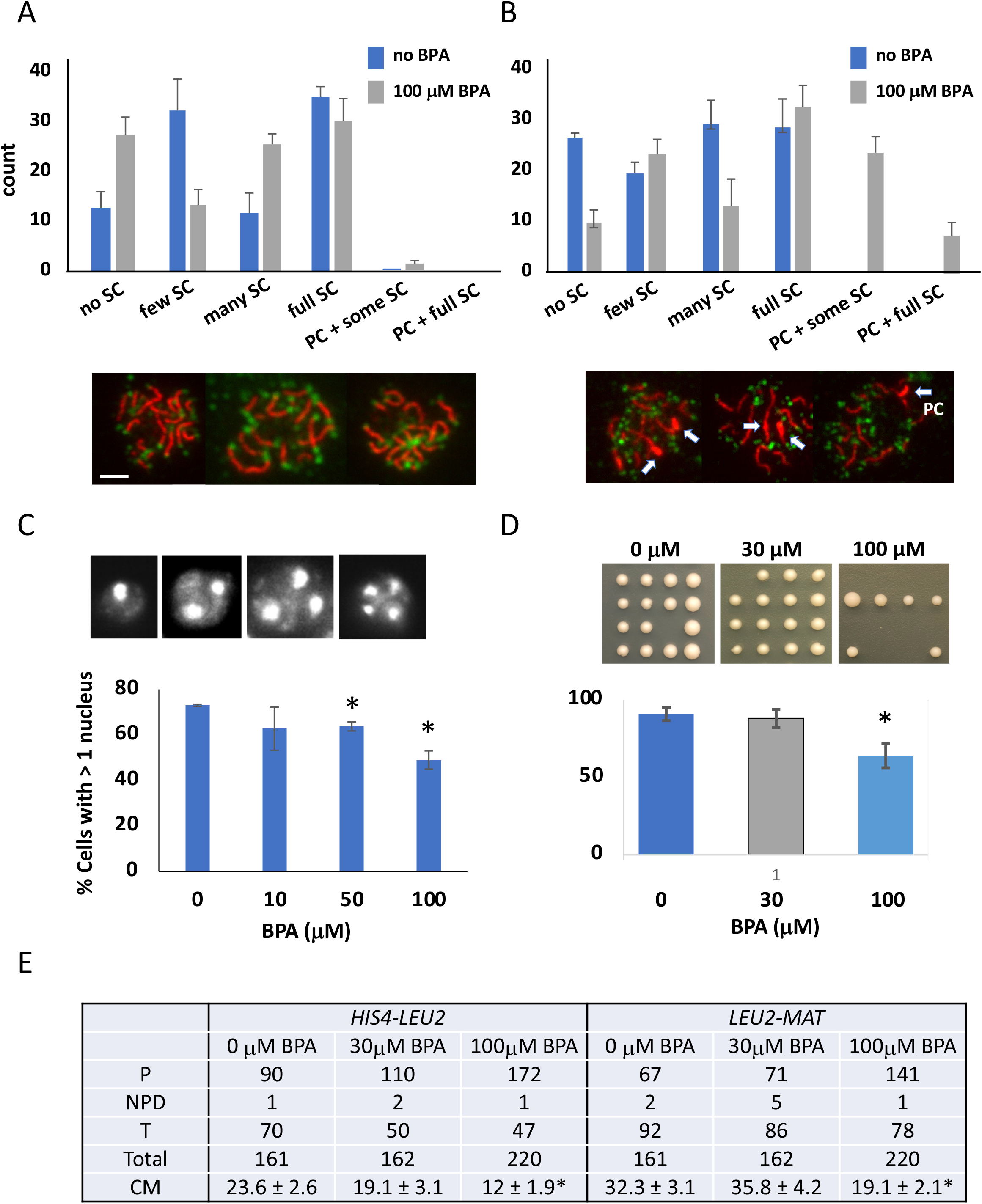
BPA affects defined aspects of meiosis in yeast. Distribution of extent of synapsis as measured by immunofluorescence staining of chromosome spreads when 100 μM BPA is added during meiosis. A) 19 hours or B) 21 hours after meiotic induction. Synapsis or the incorporation of the synaptonemal complex (SC) was detected using anti-Zip1 antibodies (red) and anti-Rap1 antibodies (green) which highlight chromosome ends. The extent of observed synapsis was classified into groups of no SC, few SC, many SC and full SC. The panels below show example chromosome spreads with full synapsis. The number of spreads having polycomplexes – aggregates of Zip1 protein that occurs when meiotic progression is delayed during prophase I – is shown. C) Panel shows the number of DAPI stained nuclei, which indicates whether meiosis I (> 2 nuclei) or meiosis II (>3 or 4 nuclei) has completed. The number of cells with > 1 nucleus indicates the gametogenesis frequency. D) Gamete viability determined by tetrad dissection. * indicates significant difference (P ≤ 0.5, t-test).

### Relative reproductive toxicity of BPA alternatives

Due to numerous studies linking BPA to reproductive toxicological effects, BPA limitations have been imposed for use in daily products (e.g. Commission Regulation (EU) 2018) resulting in the increased substitution of BPA with BPA analogs, which are not necessarily less toxic than BPA itself (Pelch et al. 2017; 2019). We therefore set out to test the relative toxicity to gametogenesis of BPA analogs by examining 19 BPA-related compounds (BADGE, BFDGE, BPAF, BPAP, BPB, BPC, BPE, 2,2’-BPF, 4,4’-BPF, BPAP, BPOPP-A, BPP, BPZ, HPP, diphenyl sulfone, hydroquinone, PHBB, TMBPA, 17 β-estradiol). **Figure 4A** shows BPA alternatives ranked in the order of meiotic toxicity using the yeast assay. Out the 19 chemicals examined, ten showed greater toxicity and nine showed lesser toxicity than BPA.

**Figure 4.**
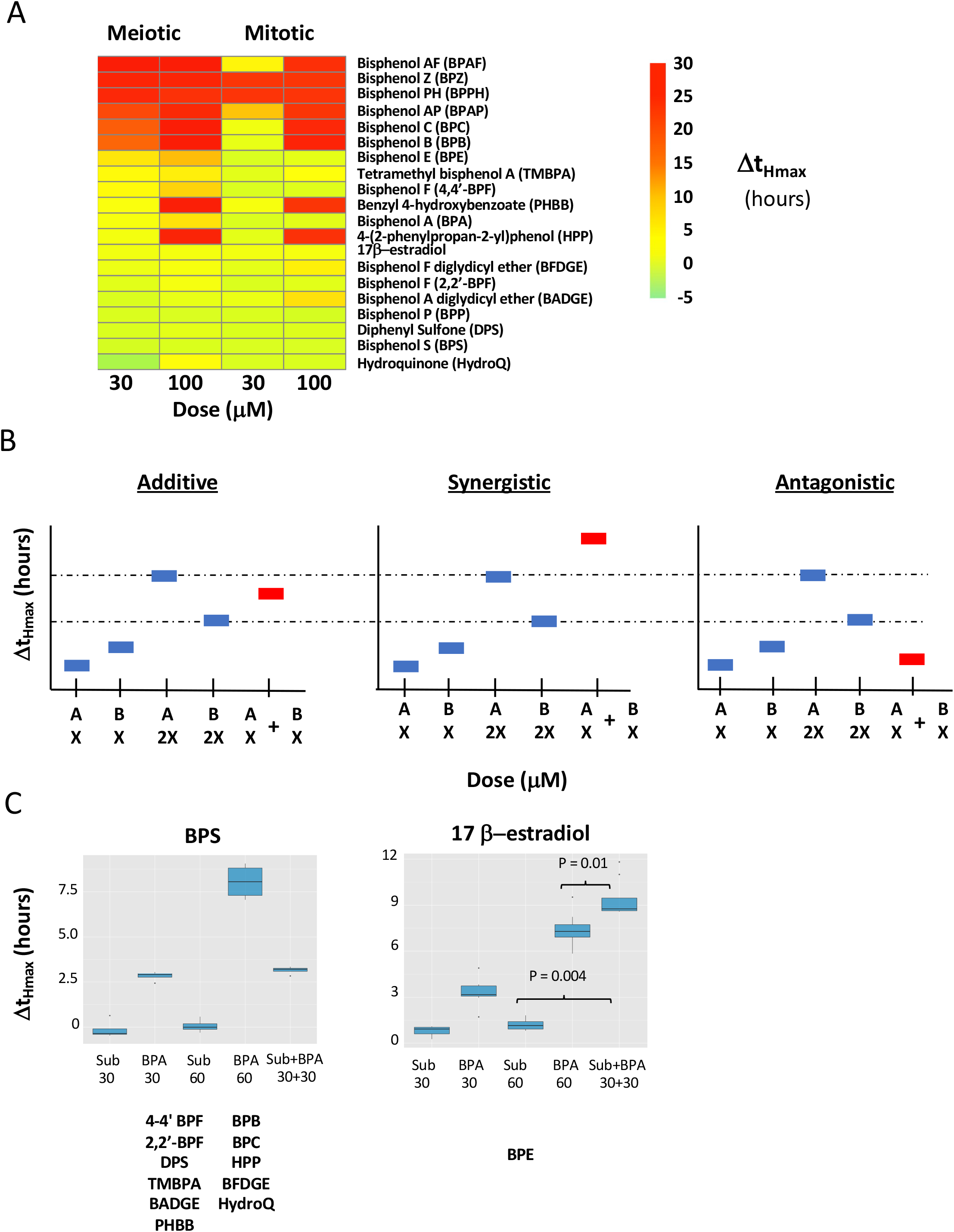
Relative reproductive toxicity of BPA alternatives. A) Heat map of Δt_Hmax_ values for BPA and 19 BPA substitutes at 30 and 100 μM doses for meiotic and mitotic assay. The chemicals are ranked first by Δt_Hmax_ 30 μM (meiotic) and then by Δt_Hmax_ at 100 μM (meiotic). B) A schematic diagram of the competition assay for BPA and its substitutes. A mixture of BPA and its substitute (sub) can show a Δt_Hmax_ equivalent to (additive), higher than (synergistic) or lower than (antagonistic) that of double the dose “X” of individual chemicals. C) Actual examples of additive (BPS) and synergistic activity (17β-estradiol) between BPA and its substitutes. Significance calculated by t-test. Additional chemicals showing additive or synergistic effects are listed under their respective examples. Graphs showing the data for each substitute with BPA can be found in Figure S3.

It is known that women are concurrently exposed to multiple potential endocrine disrupters with the potential to affect fertility (Mínguez-Alarcón et al. (2019)). The widespread adoption of BPA alternatives raises the concern that simultaneous exposure to BPA along with BPA analogs might lead to synergistic effects. To explore whether each chemical mixture acts additively or whether there are synergistic or antagonistic effects between BPA and its alternatives, we performed a series of competitive assays (see Methods) to elucidate whether such effects exist (**Figure 4B**). Out of 14 BPA substitutes, two of the BPA substitutes – BPE and 17β-estradiol, showed synergistic effects with BPA (Figure 4C, Figure S3). None showed antagonistic effects.

### Screening environmental chemicals for reproductive effects

Having demonstrated selectivity in our assay based on both known mutants and control compounds, and having shown its ability to detect meiotic defects caused by BPA, it becomes possible to apply this assay widely to measure meiotic effects of other compound classes. We thus applied our assay to an additional 179 chemicals (199 total for the entire study) (**Table S1**) spanning several environmentally relevant use categories (i.e. fire retardants, pesticides, pharmaceuticals, cosmetics, food additives, plasticizers, tobacco-related chemicals, flavorants, cleaners and industrial chemicals) and chemical classes (i.e. phthalates, per- and polyfluoroalkyl substances (PFAS), quaternary ammonium compounds (QAC), organophosphate esters (OPE)).

The majority of the chemicals were selected from a database that prioritizes chemicals for testing in order to facilitate cross comparisons of different reproductive and development assays (Abrahamsson et al. 2021). The chemicals included those suspected to negatively impact human health and those of interest to policy makers (e.g. TSCA chemicals, chemicals under consideration for EPA’s priority list). Several of these chemicals were detected in maternal and umbilical cord blood and thus are relevant to exposure during early gametogenesis (Wang 2018, 2021). Many chemicals that were toxic in worm and rodent reproductive assays were included (e.g. parathion-methyl (Narayana et al. 2006; Uzunhisarcikli et al. 2007), Bis(2-ethylhexyl) phosphate (DEHP) (Fabjan et al. 2008; Foster et al. 2001), Tris(2-chloroethyl) phosphate (TCEP) (Gulati et al. 1991), thiabendazole (Shin et al. 2019). All chemicals were barcoded to allow for blinding of the experiments. We also included seven duplicate chemicals under a different barcode as a control for measurement consistency. We performed a minimum of six replicates for each chemical at each dose (30 μM and 100 μM). All chemicals at each dose and both the positive and negative control were included in triplicate on each plate. Each plate was repeated at least once more.

We designated chemicals that showed ΔtH ? 1.5 hours (equivalent to 20% reduction in gamete viability) with a p-value ≤ 0.05 (t-test) as reproductively toxic (reprotox20). Of the total 199 compounds screened, 57 (29%) compounds were classified as reprotox20 in our assay **(Figure 5, Table S1**). We expected to find a number of compounds we classified as reprotox20 in our system given that we deliberately included many chemicals known to be reproductive toxicants in other organisms, to assess the assay’s ability to detect reproductive toxicants common across diverse organisms. We also performed a secondary HTS for diploid proliferative growth using the same chemicals, to distinguish compounds solely affecting meiosis from compounds affecting both meiosis and mitosis. For the proliferative growth assay, cells were chronically exposed throughout the assay without chemical washout. **Figure 5** illustrates chemicals ranked from highest to lowest severity, with their toxicity categorized as meiosis-specific, growth-specific, affecting both meiotic and growth. Out of the 57 reprotox20 compounds, 17 solely affected gametogenesis. Included in the top hits are 1,3 diphenylguanidine, dichlorvos, 2-phenylphenol, bisphenol E (BPE) and 1-(benzyl)quinolinium chloride, 1-dodecyl-2-pyrrolidinone, BPA and decanedioic acid and 1,10-dibutyl ester. Forty other reproductive toxicants showed toxicity for both reproductive and proliferative growth. Only ten of the compounds were designated as toxic to proliferative but not reproductive growth. The remaining 132 compounds had less or no effect in our test system (**Figure S4, Table S1**).

**Figure 5.**
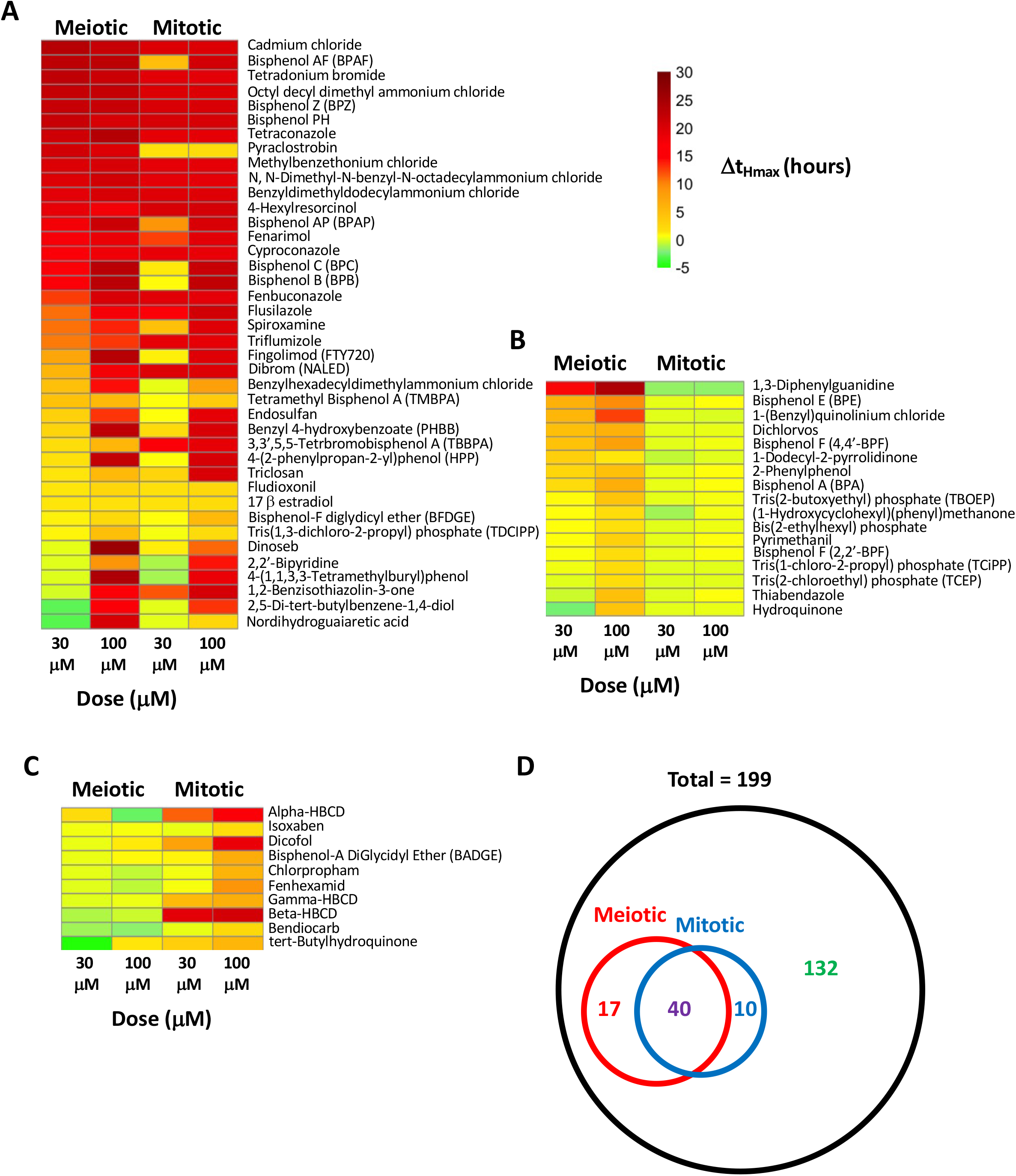
Reproductive toxicants identified by yeast HTS. 199 chemicals were evaluated for both their toxicity for meiosis and for proliferative growth (mitotic) at 30 and 100 μM doses. Chemicals were considered hits if Δt_Hmax_ > 1.5 hours and p<0.05 A) Heat map of chemicals identified as meiotic as well as mitotic hits. B) Heatmap of chemicals identified as only affecting meiosis C) Heat map of chemicals that solely affect mitotic growth. D) Venn diagram enumerating the number of hits in each category. The 132 chemicals showing neither a meiotic or mitotic effect are shown in Figure S4. Chemicals are ranked according to the meiotic Δt_Hmax_ shifts at a dose of 30 μM.

### Bisphenol and QAC chemical classes were strongest predictors for reproductive toxicity

Certain chemical classes are of interest as potential hazards to both humans and wildlife due to their persistence in the environment and potential for chemical reactivity. We thus queried whether chemicals within a particular use or chemical class were more likely to correlate with reproductive toxicity. **Figure 6** depicts the distribution of our chemicals within various usage categories (**Figure 6A**) and chemical classes (**Figure 6B**) as defined by EPA’s CompTox Chemical Dashboard (https://comptox.epa.gov/dashboard; Williams et al. 2017), an extensive searchable database that contains structure, property, toxicity, and bioassay data for collections of chemicals. In the case of usage, each compound can span many classes, however chemicals belonged to only one chemical category. We used LASSO analysis based on logistic regression as a preliminary multivariate analysis to rank the predictors (**Figure 6A, 6B;** Steyerberg et al. 2001; Vittinghoff and McCulloch 2007). Although no usage classes were linked strictly with reprotox20 toxicants in our assay, among the chemical classes we considered, bisphenol and QAC chemical structures were the strongest predictors for reproductive toxicity. Interestingly, phthalates were not found to be predictors, although this chemical category has been associated with poorer reproductive outcomes in mammals (Mesquita et al. 2021; Repouskou et al. 2021). We will discuss this apparent discrepancy below.

**Figure 6.**
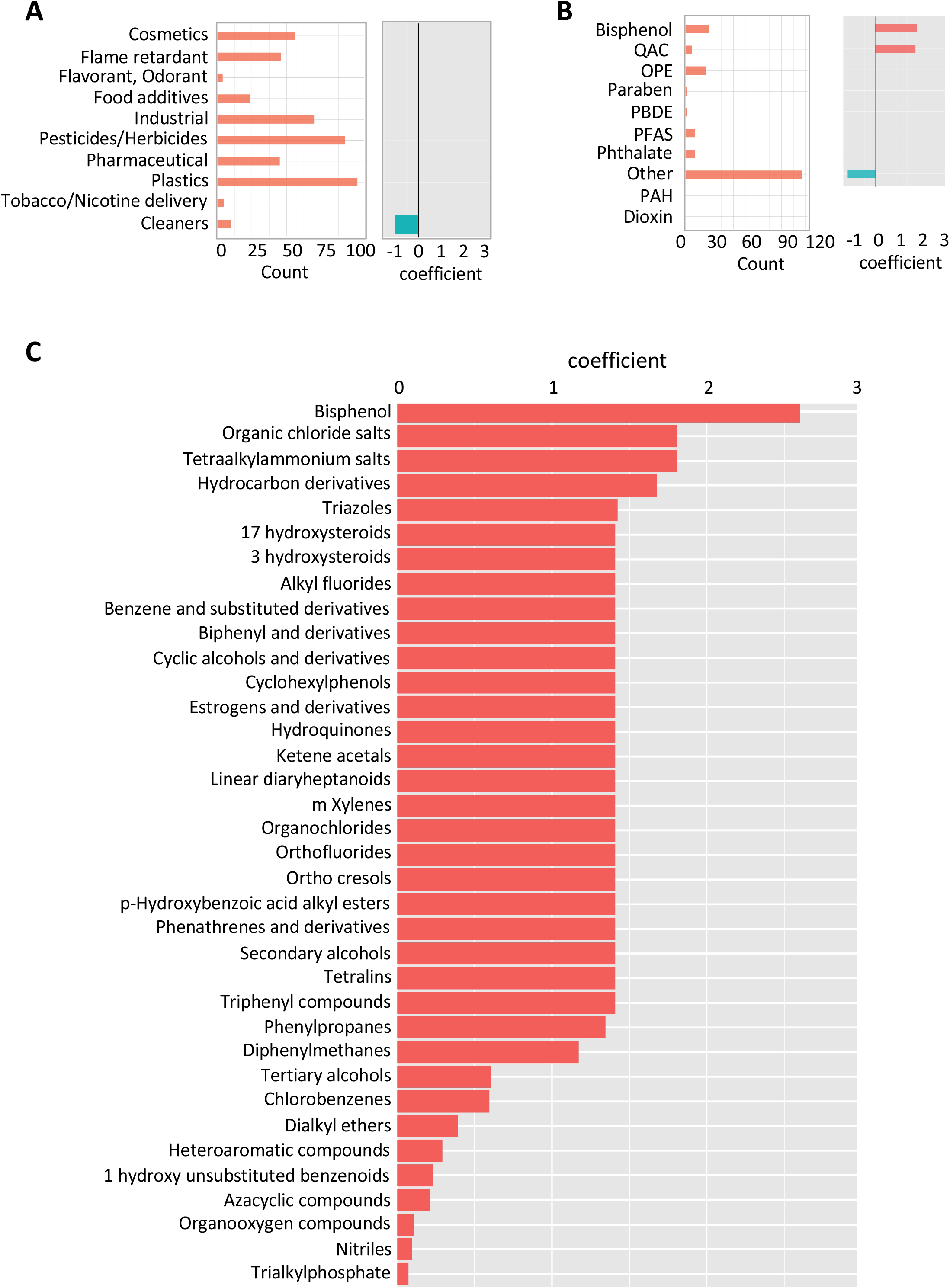
Analysis for predictors of reproductive toxicity. A) Chemical classification based on their commercial use. Plot on the left indicates the number of chemicals in each category (a chemical can have multiple uses). Chemical use categories were used in a lasso regression model to predict meiotic toxicity (lasso outcome was whether or not a chemical is a meiotic hit). The coefficients (at lambda+1SE are listed in the plot on the right. B) Similar analysis for chemical classes (non-overlapping) C) Chemical structure categories determined using the algorithm Classyfire were used in a lasso regression model to predict meiotic toxicity. Each chemical can have multiple structural elements. Only coefficients >0 are listed in the plot. The full plot can be found in Figure S5.

We also investigated which chemical structural features were more likely to associate with toxicity to reproduction. Such information will be useful towards understanding the mode of action of chemicals through possible binding partners as well as form a database from which to base algorithms used to predict the extent of toxicity such as data that informs QSAR algorithms. Chemical entities were obtained from ClassyFire, a web-based application for automated chemical structural classification (Djoumbou et al. 2016) (**Table S2**). Figures 6C shows that bisphenols, organic chloride salts, tetraalkylammonium salts and hydrocarbon derivatives were the highest predictors from the LASSO analysis. The full LASSO analysis graph is shown in **Figure S5.**

### Yeast and mammalian reproductive toxicants show significant association

One important aim of this study was to see if a yeast-based assay alone or together with other non-animal models could identify reproductive toxicants that would be relevant to mammalian gametogenesis (**Figure 7A**). Mammalian endpoints that are typically measured to evaluate reproductive toxicity include animal weight, mortality, organ weight (testes, ovaries, liver and kidney), gonadal somatic index (GSI = gonad/animal weight), sperm count, fetal adsorption, implantation success, litter size and litter viability, fetal deformation or behavioral change, mating ability and male/female sex ratio. Of these endpoints, gonad weight, GSI and sperm count are likely the most directly relevant measures of perturbations of_gametogenesis, whereas litter size, fetal adsorption and implantation success are still relevant but less direct readouts from problems occurring during gametogenesis. The remaining endpoints were considered more distal and were not included here, allowing us to focus on evaluating reproductive toxicity outcomes most relevant to gametogenesis. Comprehensive PubMed and internet searches were performed on each chemical for mammalian reproductive toxicity information (**Table S3**). If a chemical resulted in change within our criteria of a relevant reproductive endpoint (see above) the chemical was considered to show mammalian reproductive toxicity. Out of the 199 chemicals, 145 had publicly accessible data that evaluated mammalian reproductive toxicity (**Table S3**). Figure 7B illustrates that 29 chemicals scored as meiotic toxicants in the yeast assay were also deemed reproductive toxicants based on mammalian data. The association between yeast and mammalian reproductive toxicity was considered significant using a 2 x 2 contingency table (P-value ≤ 0.0001, two tailed Fisher’s exact test) (**Figure 7C**).

**Figure 7.**
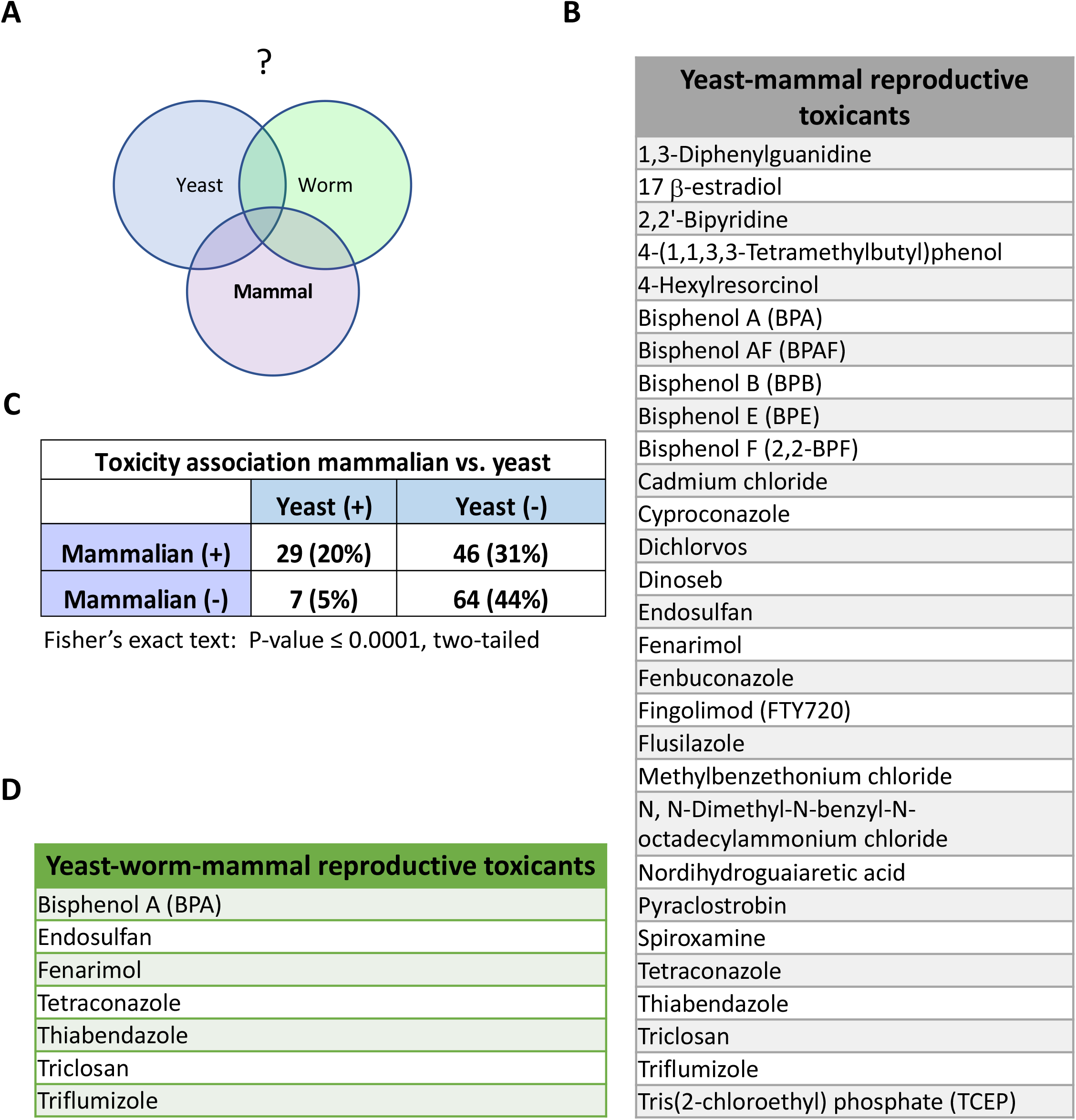
Assessment of relevance of yeast reproductive toxicants to other organisms. A) Relevance of yeast identified reproductive toxicants to mammalian gametogenesis are more probable if the same chemicals in diverse organisms overlap in their effects. B) List of chemicals that were yeast reproductive toxicants that also show mammalian reproductive toxicity based on literature search for reduction of litter size and viability, reduced gonad weights, reduced sperm count, reduced implantation and increased fetal adsorptions that could result from problems in gametogenesis. C) A Fisher’s exact test was used to evaluate whether there was any association between yeast and mammalian toxicants. D) Lists reproductive toxicants that are found in all three organisms.

The nematode *C. elegans* is one invertebrate model that has successfully be used to identify reproductive toxicants as an alternative to mammalian studies. Of 29 chemicals common between yeast and mammals, six chemicals (bisphenol A, endosulfan, fenarimol, thiabendazole, triflumizole and triclosan) were also found in the literature to be reproductively toxic in *C. elegans* (**Table S3**). Since these different organisms differ in terms of development and physiology, but share a common molecular mechanism for gametogenesis, we infer that a shared reproductive effect in all three organisms increases the likelihood that gametogenesis *per se* is affected, rather than other processes involved in reproduction.

## Discussion

In this study, we show that a yeast-based HTS is a rapid and inexpensive way to query reproductive toxicity, particularly pertaining to gametogenesis. The fact that this assay can identify BPA, which is a well-known reproductive toxicant in both mammals and other nonmammalian models (i.e. nematodes (Chen et al. 2016), zebrafish (Moreman et al. 2017)), along with the fact that BPA has similar detrimental effects to prophase I of meiosis in yeast as previously shown in worms and mammals, suggest that yeast can be useful in identifying and evaluating toxicants that impact pathways conserved in all organisms. By combining the evaluation of meiotic toxicity with an assay for proliferation, we can identify those chemicals that solely have an effect on gametogenesis suggesting that these toxicants are hitting meiosis-specific proteins and/or pathways thereby narrowing the chemical’s potential mode of action.

The yeast reproductive assay can be a preliminary step to screen through large libraries of compounds to pinpoint likely reproductive toxicants which can then be further verified in other *in vivo* systems. Multiple models are needed in order to eliminate species specific toxicities that may not be relevant to human health. Thus, combining this assay with information from other model systems will be useful for policy and regulatory purposes. Commonality between diverse organisms can provide additional evidence for human toxicity such that further examination of the effect of these chemicals on reproduction may be warranted. Our study has revealed seven chemicals: BPA, endosulfan, fenarimol, tetraconazole, thiabendazole, triflumizole and triclosan that exhibit common reproductive toxicity between yeast, nematodes and mammals (**Figure 7D**). For two of these chemicals, BPA and triclosan, human studies have been reported which show an association between higher urinary levels of these chemicals and lower antral follicle counts suggestive of reduced fecundity due to chemical exposure (Jurewicz et al. 2019; Souter et al. 2013; Zhu et al. 2019).

As shown in this study, having a rapid assay for reproductive toxicity allows quick evaluation of the relative toxicity of potential analogs and the effects of compound mixtures. In yeast, we show that the relative toxicities of some well-known BPA analogs are as follows: BPAF > BPF ~ BPA > BPS (Figure 4A). In zebrafish, hatching delay and mortality in embryos were compared for the same BPA analogs (Moreman et al. 2017) and a similar ranking of toxicities was observed: BPAF > BPA > BPF > BPS. However, in *C. elegans* it was observed that BPS is equally if not more toxic than BPA for reproductive and developmental toxicity. In other species including mammals, although BPA, BPF and BPS were compared within single studies, reproductive outcomes relating closely to gametogenesis were not evaluated (reviewed in McDonough et al. 2021).

In a previous assessment of BPA in humans that measured BPA and its primary metabolites in urine, the average concentration of BPA in humans was found to be 0.3 −0.6 μM, but with a range of 0 – 5 μM (Gerona et al. 2016). In yeast, the LOAEL or BPA concentration that gave a detectable toxic effect based on our criteria of a ΔtHmax =1.5 hours is 26 μM. This may at first be taken to suggest that the accumulated levels detected in humans are not enough to cause failure in gametogenesis. However, yeast is known to be more resistant to small molecules compared to human cells (Saunders and Rank 1982). Although we increased the chemical sensitivity of the yeast cells used in our assay by reducing the expression of the yeast multidrug transporter Pdr5, Pdr5 is not entirely eliminated, hence it remains likely that the actual concentration of any given chemical inside the yeast cell may be substantially lower than that in the surrounding media. Likewise, with respect to the human data, the concentration in urine is unlikely to be the same as the effective concentration seen by the gametes during development. It is therefore difficult to directly compare doses and sensitivities between yeast and humans.

BPA is considered a weak endocrine disruptor and many of its reproductive effects have been attributed to its role as an endocrine mimic. Although BPA has structural features compatible with binding to estrogen receptors, studies suggest that at least a part of BPA’s activity may be distinct from estradiol (Chen et al. 2019; Gould et al. 1998; Peretz et al. 2012; Ziv-Gal et al. 2013). Consistent with this notion, yeast does not have an endocrine system yet shows reproductive defects at the same stage of meiosis as seen in worms and mammals upon exposure to BPA. The synergistic effect we found between BPA and 17 β-estradiol could be suggestive that their effects may be distinct.

Phthalates are a class of chemicals found in a wide variety of consumer products due to their plasticizing abilities (Kelley et al. 2012; Net et al. 2015; Zota et al. 2014). Like BPA, phthalates have known endocrine disrupting properties and are considered reproductive toxicants (reviewed in Mesquita et al. 2021, Seda et al. 2021). In this study, out of the nine phthalates examined in yeast, none showed an effect on gametogenesis. Although this may illustrate a limitation of using yeast to detect reproductive effects of phthalates – either due to lack of an endocrine system or accessibility issues, it is interesting to note that bis(2-ethylhexyl) phosphate which shows close structural similarities to bis(2-ethylhexyl) phthalate (DEHP) does show reproductive toxicity in yeast. This opens the door to future studies to explore whether other structurally similar compounds of other phthalates can show a reproductive effect in yeast, which in turn would suggest that the structure/position of the alkyl chains of the phthalate, rather than the phthalate acid component of the molecule, is detrimental to gametogenesis. In support of this notion, Zhang et al. (2015) evaluated the structure-dependent activity of phthalate esters and phthalate monoesters binding to human androstane receptor and found that their docking results suggest that the strong binding of phthalates to the receptor arose primarily from hydrophobic interactions, π-π interactions, and steric effects of the alkyl chains.

In conclusion, this study illustrates some of the advantages of using yeast to evaluate a chemical’s impact on a complex biological process such as gametogenesis which is difficult to assess in mammals given that, at present, there is no mammalian *in vitro* culture that fully captures meiosis. In particular, due to the lack of endocrine system and the ability to directly evaluate meiosis, yeast allows us to distinguish an effect on gametogenesis from other reproductive perturbations. This study not only offers insights on the impact of these chemicals on yeast gametogenesis, but also provides a strategy for the prioritization of these chemicals for further study of reproductive toxicity in rodent models and in humans. Moreover, the ability to evaluate greater numbers of chemicals using the yeast model can provide data for QSAR modeling that can reveal relationships between structural properties of chemical compounds and biological activities and aid the development of predictive algorithms for reproductive toxicity. Target identification can also be pursued via use of commercial mutant libraries spanning the yeast genome as means to elucidate the genes and biological pathways that contribute to the observed toxicity.

## Acknowledgements

The authors are grateful to Wallace F. Marshall, Julia Varshavsky and Juleen Lam for critical reading of the manuscript and for John Boscardin for statistical consultation. This work was supported by grants from the National Institutes of Health (R01ES027051 02S1 – JCF, PA, TJW; R01GM137126 – JCF)

---

**Figure S1.**
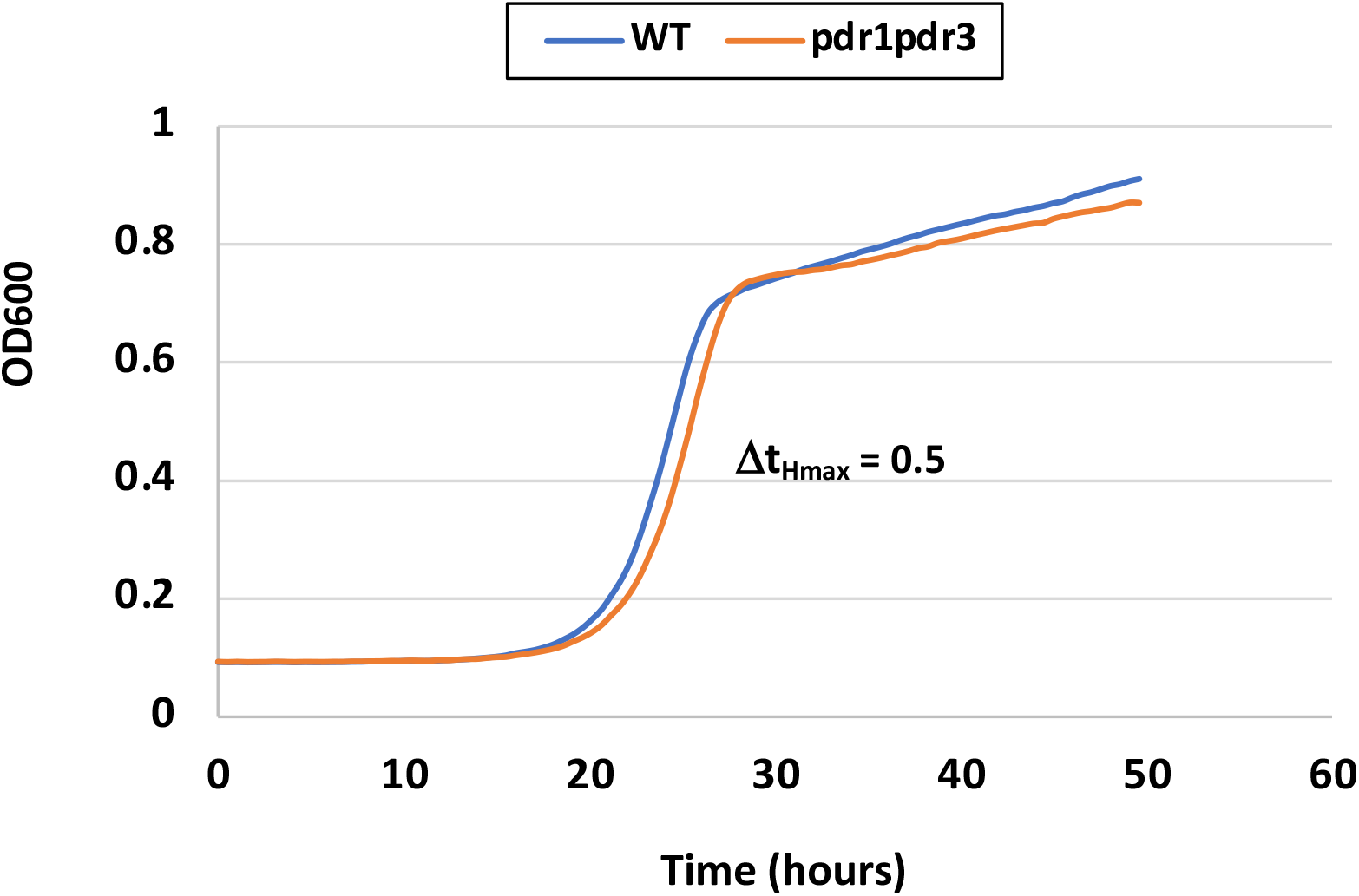
Minimal effect of the *pdr1Δ pdr3Δ* double mutant on gametogenesis. A non-statistically significant shift of 0.5 hrs is seen in the *pdr1Δ pdr3Δ* double mutant as compared to wildtype.

**Figure S2.**
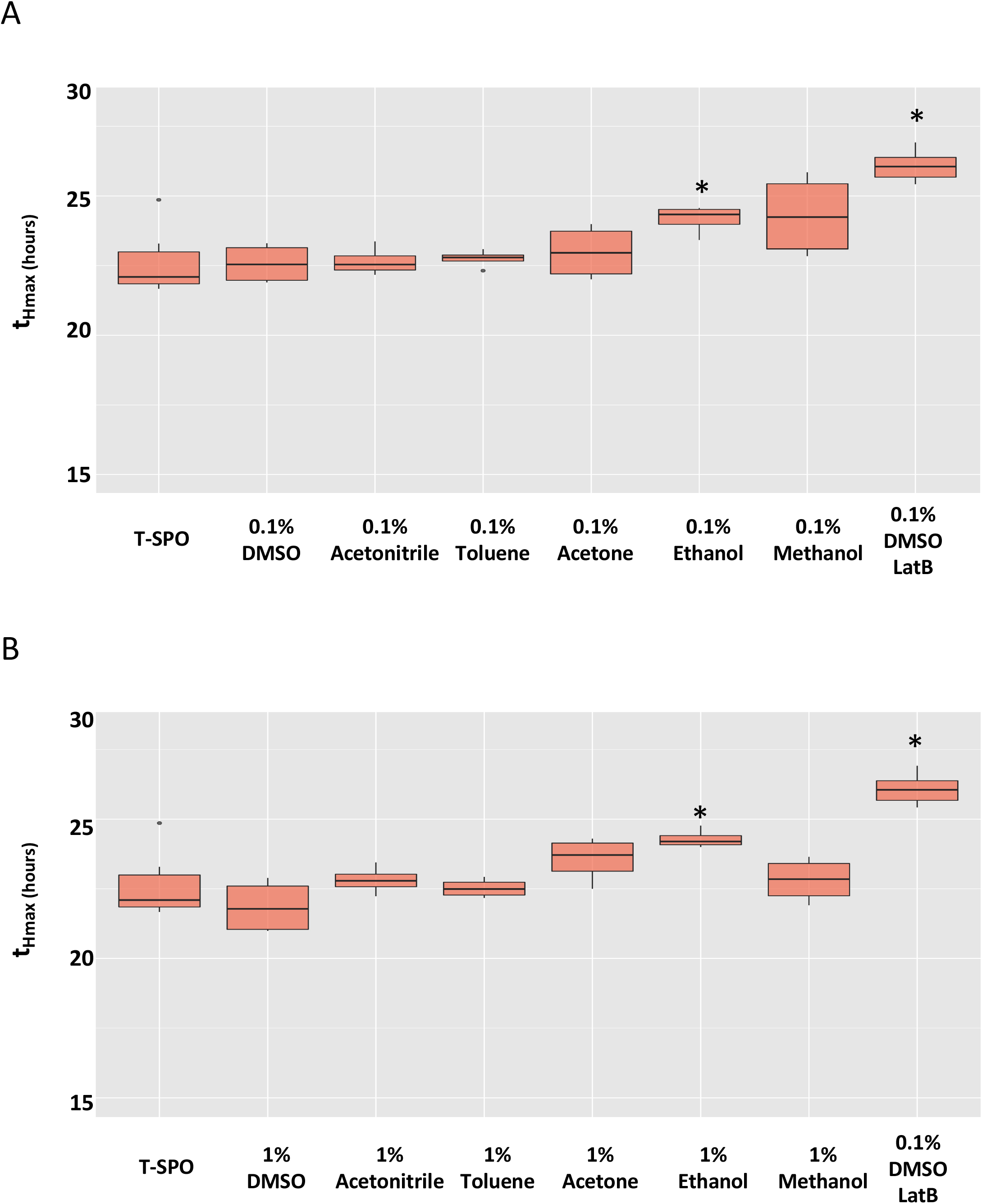
Time to halfmax using various solvents in T-SPO. Several solvents were compared for toxic effects to gametogenesis. A) 0.1% solvent comparison. B) 1% solvent comparison. T-SPO is the meiotic media used to induce gametogenesis. 0.5 μM Latrunculin B (LatB) in 0.1% DMSO is used as a positive control. * Indicates significant difference P<0.05, t-test.

**Figure S3.**
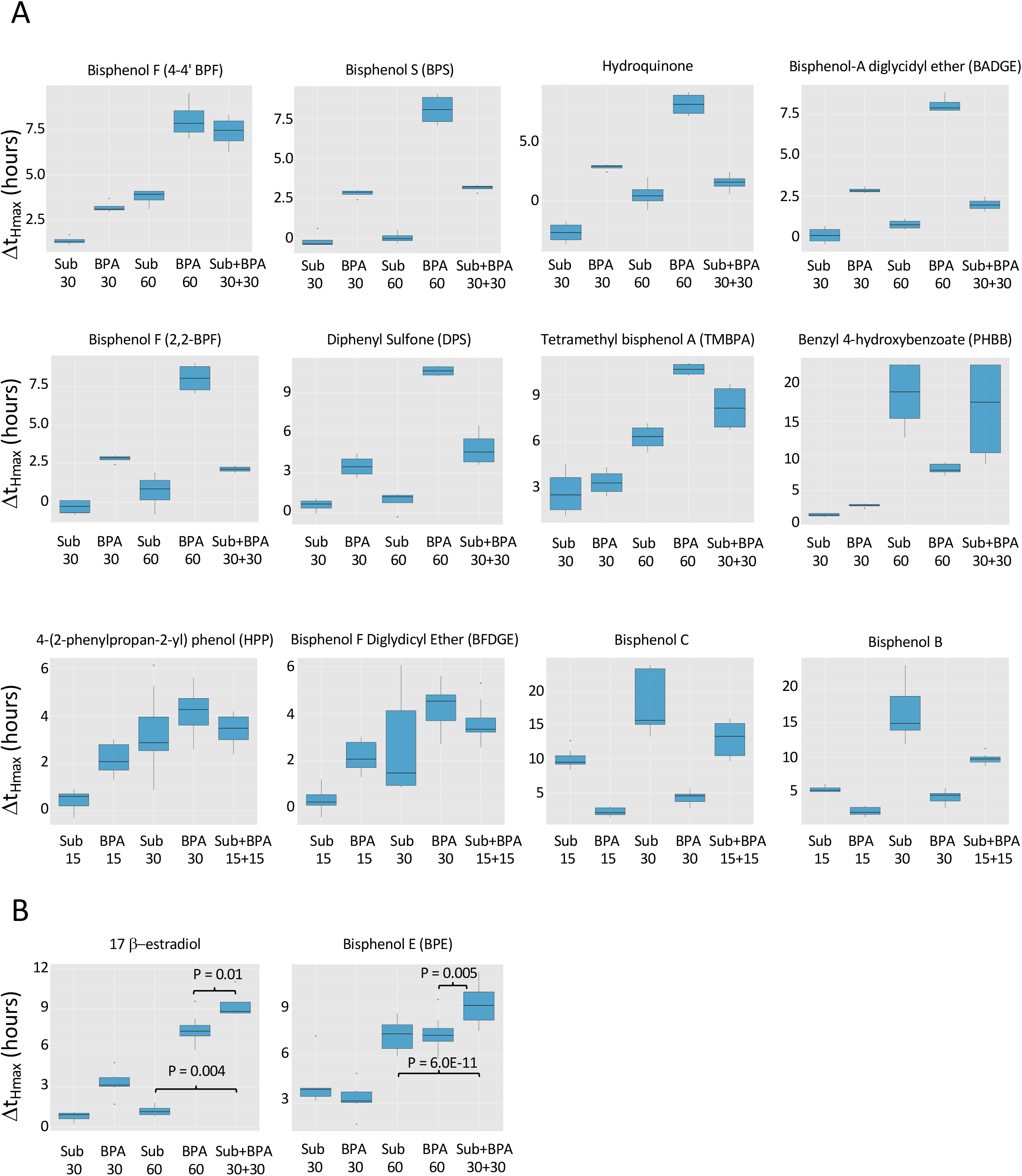
Competitive Assays for Additive, Synergistic and Antagonistic Effects for BPA vs. BPA substitutes. Sub refers to the BPA substitute in the heading used in the assay. The numbers in the x-axis indicate doses used.

**Figure S4.**
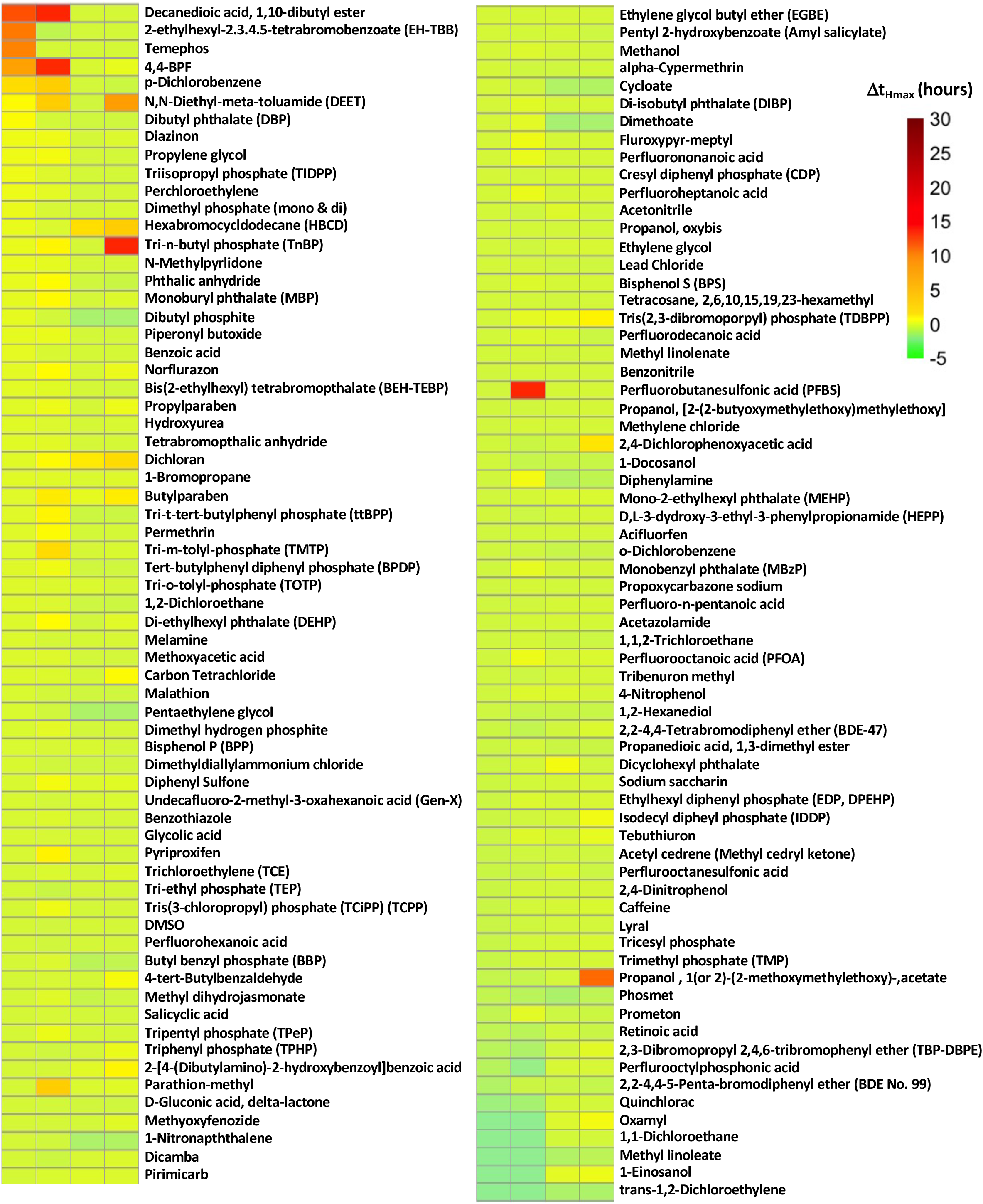
Heat map of chemicals that did not show any significant halfmax shift in both meiotic and mitotic assay. (p>0.05 or Δt_Hmax_ <1.5). Ranked by 30μM meiotic shifts.

**Figure S5.**
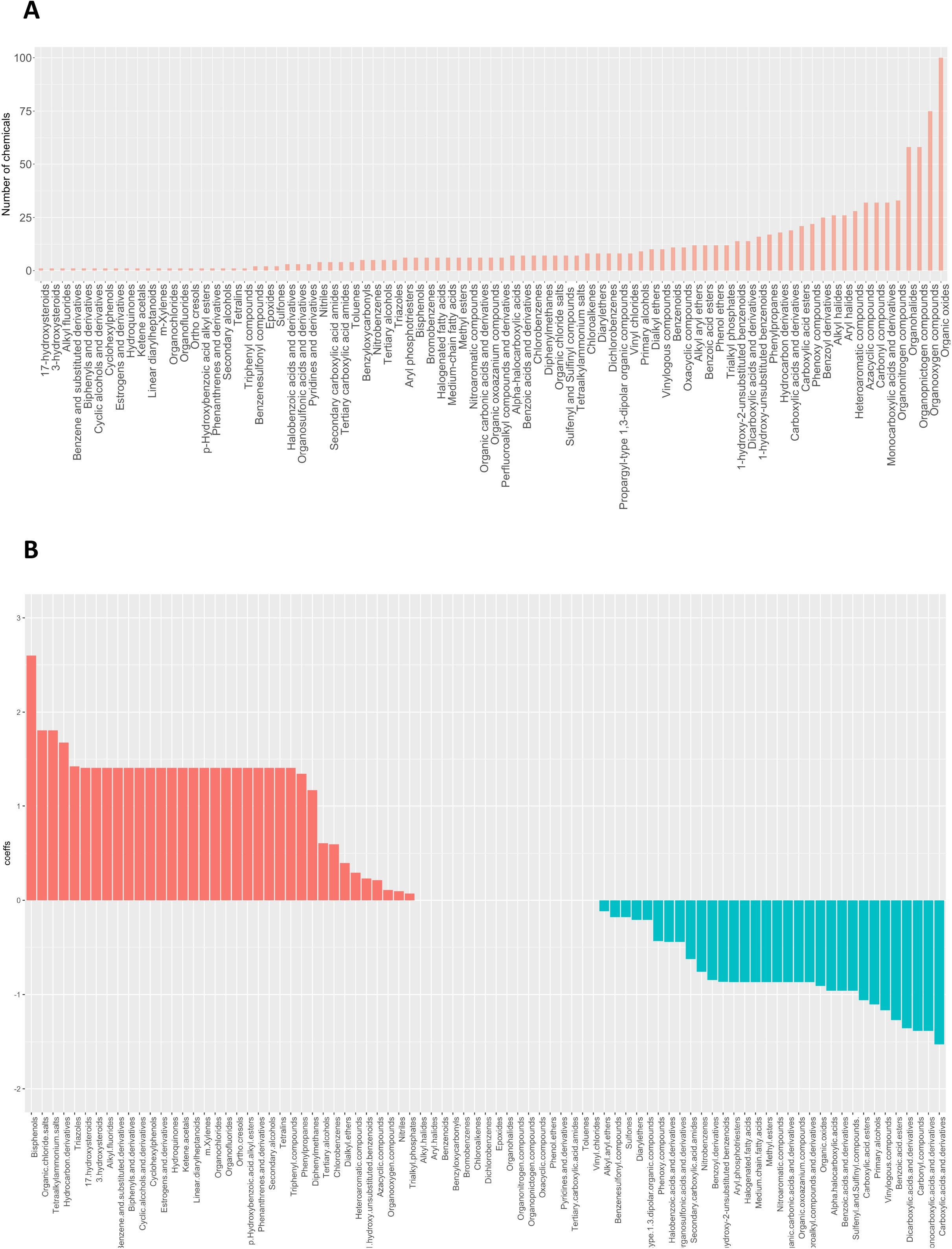
Complete chemical structures and Lasso regression. A) Chemical structural components and their prevalence as determined by the Classyfire algorithm. B) Chemical structural components were used in a lasso regression model to predict meiotic toxicity

